# HIF-1α Contributes to the Progression of Chronic Obstructive Pulmonary Disease

**DOI:** 10.1101/2022.01.09.472256

**Authors:** Kedong Zhang, Feng Zhou, Caixia Zhu, Liang Yuan, Defu Li, Jian Wang, Wenju Lu

## Abstract

**Background:** Hypoxia-inducible factor-1α (HIF-1α) plays an important regulatory role in inflammatory and hypoxic diseases. Higher HIF-1α level was found in lungs of chronic obstructive pulmonary disease (COPD) patients, however, its role in cigarette smoke (CS)-induced COPD has not been fully studied. Digoxin has been showed to inhibit HIF-1α translation and block HIF-1α activity and thus is often used as the HIF-1α inhibitor. Therefore, in the present study, we chose digoxin as the inhibitor to investigate whether HIF-1α contributes to the progression of COPD and possible mechanism.

**Methods:** CS-exposed mice were intragastrically treated with different doses of digoxin, and COPD-associated phenotypes such as pathological changes in lungs, inflammation, lung function and mucus secretion in airways were evaluated. Meanwhile, CSE-treated A549 cells were administrated with digoxin or S7959. Moreover, EMT-associated markers together with HIF-1α\TGF-β1\Smad3 signaling pathway were detected both *in vivo* and *in vitro*.

**Results:** The level of HIF-1α was significantly increased in lungs of COPD mice and CSE-exposed A549 cells, which was markedly suppressed by digoxin. Moreover, digoxin inhibited CS-induced inflammatory responses, lung function decline, and mucus hyper-secretion in COPD mouse model. In *in vitro* studies, digoxin decreased CSE-induced pro-inflammatory cytokine release. Importantly, CS-induced or CSE-induced EMT and up-regulation of HIF-1α/TGF-β1/Smad pathway was inhibited by digoxin. Additionally, S7959 mitigated CSE-induced EMT in A549 cells.

**Conclusions:** Digoxin can protect CS-induced COPD and prevent CS-induced EMT possibly through HIF-1α/TGF-β1/Smad3 signaling pathway. This study suggests HIF1-α could be a potential intervention target for COPD prevention and treatment, especially for EMT in CS-induced COPD.

## 1. Introduction

Chronic obstructive pulmonary disease (COPD), a progressive lung disease, is most caused by long-term smoke (1) and predicted to be the third leading cause of death globally by 2020 year (2). Its pathological changes mainly include emphysema, respiratory structural remodeling, and airway goblet cell hyperplasia and mucus hyper-secretion (3). The commonly-used medications for COPD are mainly bronchodilators and inhaled corticosteroids (2), but they cannot slow down lung function decline and improve the prognosis (2). Therefore, there is an enormous need for us to find new intervention target or develop new drugs to improve the progression and prognosis of COPD.

Hypoxia-inducible factor-1α (HIF-1α), a subunit of heterodimeric transcription factor HIF-1, activates the transcription of target genes under hypoxia conditions (4). Moreover, it is an important regulator of cellular responses to inflammation and oxidants (5). Inflammation and oxidative stress are important pathogenesis for COPD (2). Some studies also reported that HIF-1α might promote inflammatory responses (6, 7), airway goblet cell hyperplasia (8), and emphysema (9) in COPD. However, its role in cigarette smoke (CS)-induced COPD has not been fully studied, especially in COPD-related epithelial-mesenchymal transition (EMT).

EMT of airway epithelium is increased in COPD patients and normal smokers, and it contributes greatly to the occurrence and progression of COPD (10–13). Moreover, EMT is involved in the respiratory structural remodeling and airway fibrosis in COPD (14). HIF-1α plays a key role in EMT of renal fibrosis and mammary cancer cells (15, 16). But so far, the role of HIF-1α on EMT in CS-induced COPD has not been reported. Transforming growth factor-β1 (TGF-β1)/Smad pathway plays a crucial role in triggering EMT (17). Phosphorylated Smad2 and Smad3 form a stable complex with Smad4 and then transfer into the nucleus, subsequently modulating the transcription of EMT-associated genes (18). Besides, HIF-1α protein synthesis can regulate the activation of TGF-β1/Smad signaling pathway (19). Therefore, HIF-1α might contribute to the progression of EMT in COPD through the TGF-β1/Smad signaling pathway.

Digoxin has been showed to inhibit HIF-1α protein translation and block HIF-1 activity and thus is often used as a HIF-1α inhibitor (20, 21). Therefore, in the present study, we investigated the effects of HIF-1α on CS-induced COPD by administrating the CS-exposed mice or cigarette smoke extract (CSE)-stimulated A549 cells with digoxin. Furthermore, we determined whether HIF-1α contributed to the progression of EMT in COPD and whether this effect was mediated through the TGF-β1/Smad signaling pathway.

## 2. Materials and Methods

### 2.1 Animals and experiment design

Male C57BL/6 mice, weighting 18-20 g, were purchased from Guangdong Medical Laboratory Animal Center (Guangdong, China), and were allowed free access to food and water in a restricted specific pathogen-free room with controlled temperature (25°C) under 12 h light/12 h dark cycle. All animal experiments were performed according to the Criteria of the Medical Laboratory Animal Administrative Committee of Guangdong and the Guide for Care and Use of Laboratory Animals of Guangzhou Medical University. And the protocols were approved by Ethic Committee for Experiment Research, the First Affiliated Hospital, Guangzhou Medical University. The mice were randomly divided into five groups: control group, CS group, CS plus digoxin groups (0.02mg/kg and 0.1mg/kg) and digoxin (0.02mg/kg) group. To establish the COPD mouse model, mice were exposed to CS that were produced by 9 filtered cigarettes (Plum brand, Guangdong Tobacco Industry Co., Ltd., Guangdong, China) for 4 h per day, 6 days per week, 24 weeks together in a whole-body exposure chamber. After 16-week CS exposure, mice in the CS plus digoxin groups were intragastrically treated with different doses of digoxin (Sigma, D6003) once a day, 6 days a week, 8 weeks together before exposure to CS. Meanwhile, mice in the CTL and CS groups were given an equal amount of CMC-Na (0.5%), which as a suspending agent for digoxin. After 8-week digoxin or CMC-Na administration, all mice were sacrificed and used to study the effects of digoxin on COPD.

### 2.2 Cell culture and treatment

A549 cells were cultured in DMEM containing 10% fetal bovine serum, 100 mg/L penicillin, and 100 mg/L streptomycin in a humidified incubator at 37◻with 5% (v/v) CO_2_. When cell abundance reached 60%-70%, A549 cells were treated with digoxin or S7959 (a Smad3 inhibitor) for 2 h before 48-h CSE (2%) treatment. Each cell experiment was repeated five times. Digoxin or S7959 was dissolved in DMSO at a concentration of 50nM or 100μM. CSE was freshly prepared from Plum brand filtered cigarettes within 30 min prior to CSE treatment according to a described protocol (22). And prepared fresh CSE was regarded as 100%. All other culture reagents used in our study were purchased from Gibco (Carlsbad, CA, United States).

### 2.3 Assessment of lung function

Pulmonary function was evaluated just as previously described (23). The total lung capacity (TLC), functional residual capacity (FRC), forced vital capacity (FVC), resistance index (RI) and forced expiration volume at 50 ms (FEV_50_) were obtained according to the Buxco resistance/compliance application manual.

### 2.4 Western blot analysis

Western blot analysis was performed as described by Li et al. (23). After PVDF membranes were incubated with horseradish peroxidase (HRP)-conjugated anti-mouse or anti-rabbit secondary antibodies, western blotting images were obtained by Tanon 5200 chemiluminescence imaging system (Shanghai Tanon Science & Technology, Shanghai, China). Semi-quantitative analyses of immunoblots were performed by using the Image J. The primary antibodies used in our study were as follows: mouse-anti-β-actin antibody (sc-47778, Santa Cruz Biotechnology, Dallas, TX), rabbit-anti-HIF-1α antibody (NB100479, Novus Biologicals, Littleton, Colorado, USA), mouse anti-Ecadherin antibody (Cell Signaling Technology, Danvers, CA, United States), rabbit anti-Vimentin antibody (Cell Signaling Technology, Danvers, CA, United States), rabbit anti-ZO-1antibody (RA231621, Cell Signaling Technology, Danvers, CA, United States), rabbit anti-phospho-Smad3 antibody (9523S, Cell Signaling Technology, Danvers, CA, United States), and rabbit-anti-Smad3 (9520T, Cell Signaling Technology, Danvers, CA, United States). β-actin was used as the internal control.

### 2.5 Real-time PCR analysis

Total RNA was extracted from lung tissues or A549 cells using Trizol rea gent (Invitrogen) and reversely transcribed into cDNA using the PrimeScript RT reagent Kit with gDNA Eraser (TAKARA, Japan). Primer sequences for target genes (HIF-1α, TGF-β1, Smad3, IL-6, IL-1β and TNF-α) were as follows: Mouse HIF-1α (Fwd 5’-GATGACGGCGACATGGTTTAC-3’ and Rev 5’-CTCACT GGGCCATTTCTGTGT-3’); Mouse TGF-β1 (Fwd 5’-ATCTCGATTTTTACCCTG GTGGT-3’ and Rev 5’-CTCCCAAGGAAAGGTAGGTGATAGT-3’); Mouse Sma d3 (Fwd 5’-AGATGACAGTGCAGCAGTGGGT-3’ and Rev 5’-CAGCAGAGGA GAAGGGGTAAAGAG-3’); Mouse IL-1β (Fwd 5’-GCCTCGTGCTGTCGGACC CATAT-3’ and Rev 5’-TCCTTTGAGGCCCAAGGCCACA-3’); Mouse IL-6 (Fw d 5’-TCACAGAAGGAGGGCTAAGGACC-3’and Rev 5’-TCACAGAAGGAGTG GCTAAGGACC-3’); Mouse TNF-α (Fwd 5’-CCCTCCTGGCCAACGGCATG-3’ and Rev 5’-TCGGGGCAGCCTTGTCCCTT-3’); mouse 18S (Fwd 5’-GCA ATT ATT CCC CAT GAA CG-3’and Rev 5’-GGC CTC ACT AAA CCA TCC AA-3’); Human HIF-1α (Fwd 5’-TGCTTGGTGCTGATTTGTGAACC-3’ and Rev 5’-CTGTCCTGTGGTGACTTGTCC-3’); Human TGF-β1 (Fwd 5’-AAGGACCTC GGCTGGAAGTG-3’ and Rev 5’-CCGGGTTATGCTGGTTGTA-3’); Human Sma d3 (Fwd 5’-GAGTGAAGATGGAGAAACCAGTGAC-3’ and Rev 5’-GTAGTAG GAGATGGAGCACCAGAAG-3’); Human GAPDH (Fwd 5’-TGACTTCAACAG CGACACCCA-3’ and Rev 5’-CACCCTGTTGCTGTAGCCAAA-3’). 18S and GAPDH were used as an internal control in mouse lung tissue and A549 cells, respectively. And all the primers were synthesized by Sangon Biotech (Shanghai, China)

### 2.6 Bronchial alveolar lavage fluid (BALF) analysis

Mice were sacrificed with 1% pentobarbital sodium (50 mg/kg, i.p.). Then the lungs underwent lavage with 0.6 ml saline for three times. BALF was collected and centrifuged (800 g, 5 min, 4 ◻). Then the sediment cells were resuspended with 1ml saline for cell classification and counting. The cells were subjected to Giemsa staining for differential counting of neutrophils, macrophages and lymphocytes. The supernatant was stored at −80°C for pro-inflammatory cytokine detection by ELISA.

### 2.7 Lung pathological analysis

Left lungs were perfused with 10% buffered formalin and then immediately immersed and fixed in this fixative solution for 24 h. The lung tissues were embedded in paraffin, cut into 3-mm-thicksections and then stained with hematoxylin and eosin (H&E) and periodic acid-Schiff (PAS, Shanghai Sun Biotechnology, Shanghai, China) for histological examination.

Goblet cells were identified by PAS staining. PAS-positive cells were quantified by a previously-described semi-quantitative method with slight modification(24). Briefly, PAS-positive and total epithelial areas were measured by Image Pro-Plus, and the goblet cell metaplasia score was calculated as the percentage of the PAS-stained area to the total epithelial area after scoring 3 or 4 different areas per slide from 5 slides in each treatment group. Moreover, the average alveolar intercept of the lungs was measured by Image Pro-Plus.

### 2.8 Enzyme-linked immunosorbent assay (ELISA)

BALF and cell supernatant were prepared for ELISA. The concentration of tumor necrosis factor-α (TNF-α), interleukin-6 (IL-6), TGF-β1 and monocyte chemotactic protein-1 (MCP-1) was measured using the ELISA kit following the manufacturer’s protocol (eBioscience Affymetrix, Santa Clara,CA, United States). The levels of Muc5ac and Muc5b in BALF were also measured according to Lu et al. (25).

### 2.9 Measurement of body weight and liver, spleen, kidney indices

All mice were weighed once a week during the first 16 weeks of CS exposure and twice a week during digoxin treatment. In addition, the liver, spleen, and kidney were blotted dry and weighed at the terminal of CS exposure. The ratios of liver, spleen and kidney weight to the body weight were counted as liver, spleen and kidney indices, respectively.

### 2.10 Hematocrit (HCT) measurement

Briefly, at the terminal of chronic CS exposure, whole blood was collected into capillary tubes (0.5 mm outside diameter, VWR Scientific, Radnor, PA) via right ventricle puncture with K2EDTA as an anticoagulant and centrifuged at 7,000 rpm for 5 min, and read on a hematocrit chart (VWR Scientific).

### 2.11 Data statistics

Statistical analyses were performed using SPSS 16.0. One-way ANOVA followed by Bonferroni post-hoc test was used for comparisons between more than two groups. Data were presented as mean ± SEM, and difference with p-value <0.05 was regarded significant.

## 3. Results

### 3.1 Digoxin protects against CS-induced lung function decline and hypoxia in mice

In order to investigate the potential role of digoxin as a HIF-1α inhibitor in CS-induced COPD, COPD mice induced by six-month consecutive CS exposure were treated with two doses of digoxin (0.02 mg/kg and 0.1 mg/kg). CS-exposed mice exhibited obvious lung function decline (**Fig. 1A-E**), manifested by a decrease in FEV_50_/FVC and higher values in TLC, FRC, FVC and RI compared with CTL group. Hematocrit, an indicator of chronic hypoxia, was increased in mice exposed to CS compared with CTL group (**Fig. 1F**). The aforementioned COPD-like symptoms, including lung function decline and chronic hypoxia, were ameliorated by digoxin (mainly 0.1 mg/kg). These results demonstrate that digoxin protects against lung function decline and hypoxic changes in the COPD mouse model.

**Figure 1.**
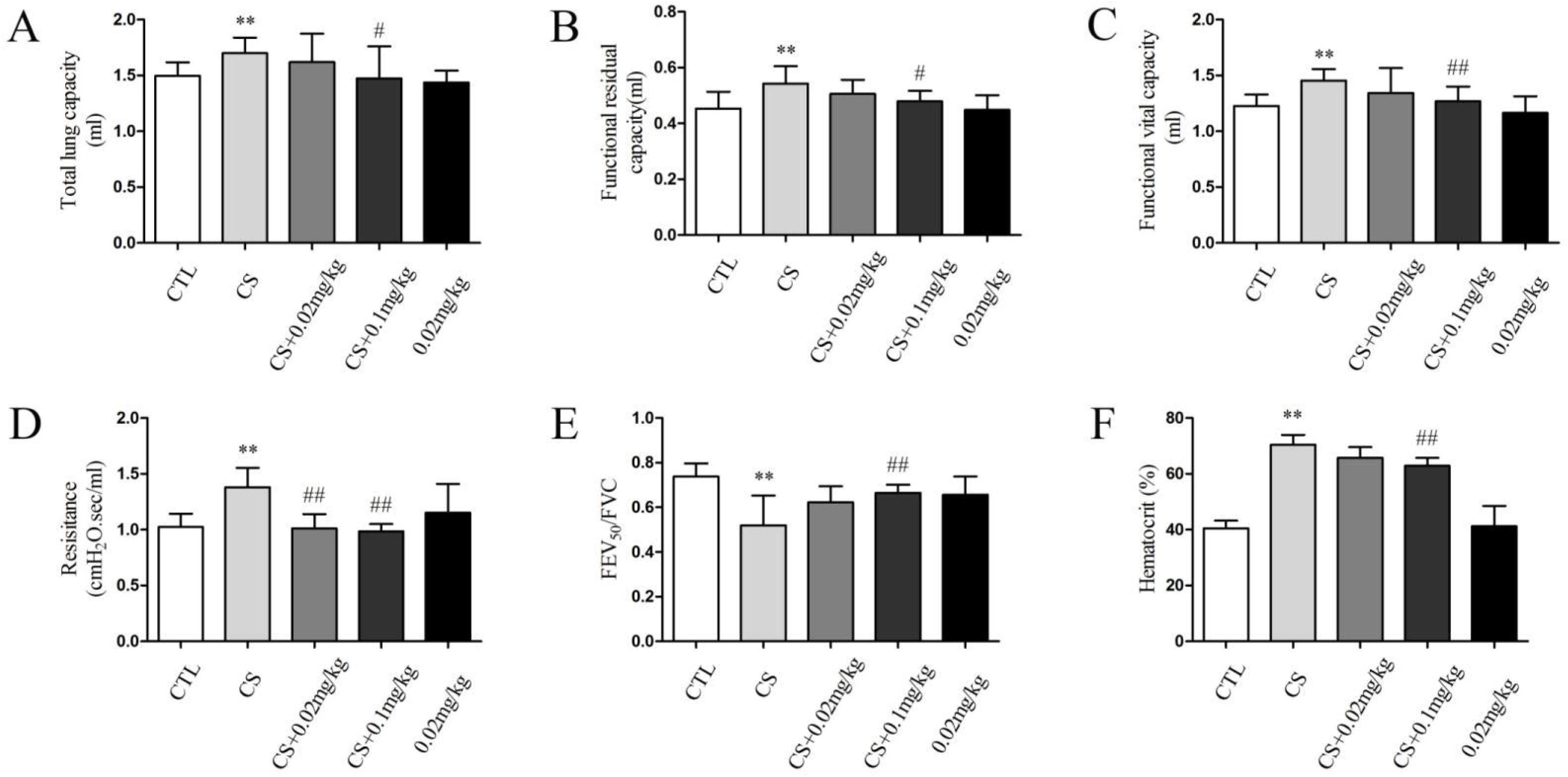
Digoxin protects against CS-induced lung function decline and hypoxia in mice. To investigate the role of HIF-1α on CS-induced COPD, mice exposed to CS for 6 months were intragastrically treated with digoxin (0.02 and 0.1 mg/kg). (A) TLC, (B) FRC, (C) FVC, (D) Resistance, (E) FEV_50_/FVC and (F) Hematocrit were evaluated. All data were shown as the mean ± SD; n= 8-10 per group. Statistical significance was assessed by one-way ANOVA. \**p< 0.05* and ^**^*p< 0.01* versus control group; ^#^*p< 0.05* and ^##^*p< 0.01* versus CS group.

### 3.2 Digoxin decreases CS-induced or CSE-induced inflammation in both mice and A549 cells

Long-term smoking can cause chronic inflammation in lungs, which is a contributing factor in COPD pathogenesis (26). Digoxin attenuated CS-induced increases in the number of total inflammatory cells and macrophages in BALF (**Fig. 2A-B)**. Moreover, the mRNA levels of TNF-α, IL-6 and IL-1β in lung tissues (**Fig. 2C-E)** as well as the TNF-α, IL-6 and MCP-1 protein levels in BALF (**Fig. 2F-H)** of CS-exposed mice were reduced by digoxin treatment (both 0.02 and 0.1 mg/kg,). To extend these findings to a human system, A549 cells were exposed to 2% CSE for 48 h in the presence or absence of digoxin (50 nM), and then pro-inflammatory cytokine levels in culture supernatant were measured. Consistently, digoxin treatment significantly reduced CSE-induced IL-6 and IL-8 release in A549 cells **(Fig. 3I-J)**. Taken together, these results suggest that digoxin reduces CS-induced or CSE-induced inflammatory responses in both mouse lungs and A549 cells.

**Figure 2.**
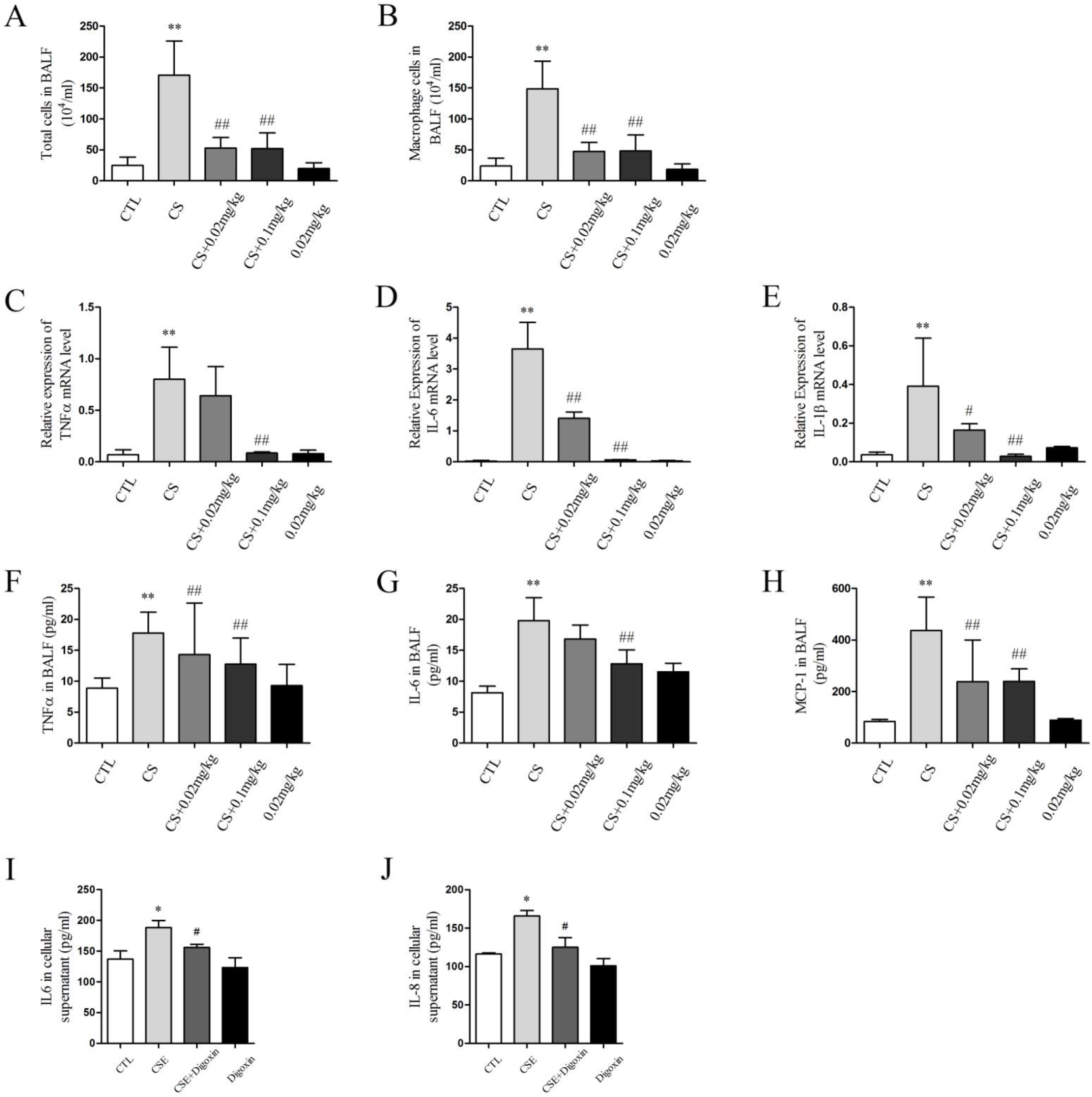
Digoxin decreases CS-induced or CSE-induced inflammation in both mice and A549 cells. (A-B) The number of total inflammatory cells and macrophages in BALF were measured by cell counting. The mRNA levels of TNF-α, IL-6, and IL-1β in mouse lung tissues were measured by real-time PCR (C-E). The protein levels of TNF-α, IL-6 and MCP-1 in BALF (F-H) were measured by ELISA. To further explore the effect of HIF-1α on CSE-induced inflammatory responses *in vitro*, A549 cells were treated with digoxin for 1 h before 48-h CSE exposure. IL-6 and IL-8 levels in cell supernatant (I-J) were measure by ELISA. All data were shown as the mean ± SD; n= 8-10 per group in mice, n=5 per group in A549 cells. Statistical significance was assessed by one-way ANOVA. \**p< 0.05* and ^**^*p< 0.01* versus control group; ^#^*p< 0.05* and ^##^*p< 0.01* versus CS group.

**Figure 3.**
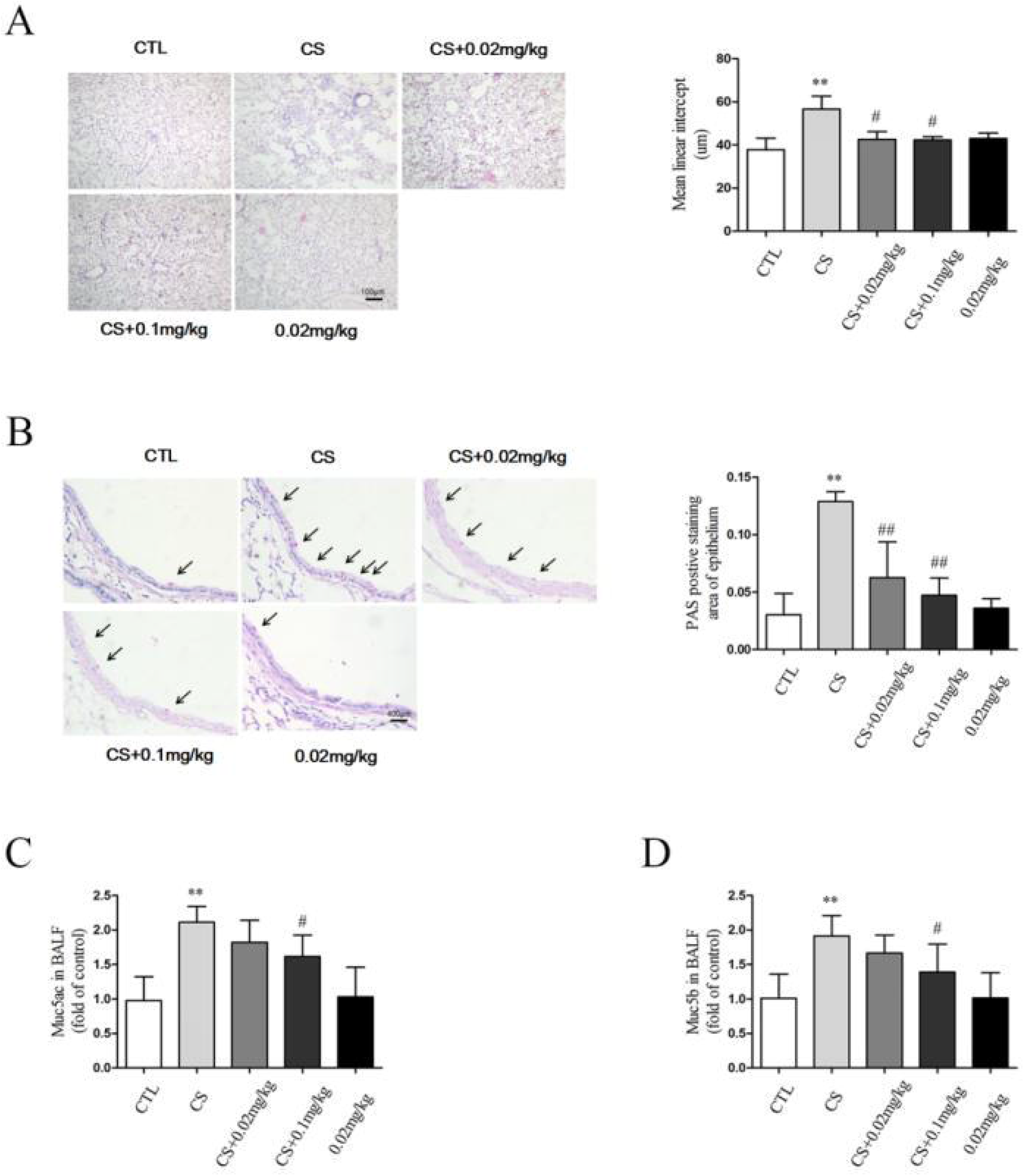
Digoxin attenuates CS-induced emphysema, airway goblet cell hypertrophy and hyperplasia, and airway mucus hyper-secretion in mice. Lung tissue sections were subjected to H&E and PAS staining. (A) The mean linear intercept (×100 magnification) and (B) goblet cell hyperplasia score (×400 magnification) in mouse lungs were analyzed using Image-Pro Plus 6.0 software. The protein levels of Muc5ac and Muc5b in BALF were measure by ELISA (C-D). All data were shown as the mean ± SD; n= 8-10 per group in mice. Statistical significance was assessed by one-way ANOVA. \**p< 0.05* and ^**^*p< 0.01* versus control group; ^#^*p< 0.05* and ^##^*p< 0.01* versus CS group.

### 3.3 Digoxin attenuates CS-induced emphysema, airway goblet cell hypertrophy and hyperplasia, and airway mucus hyper-secretion in mice

Airspace enlargement resulting from alveolar destruction was found in lungs of CS-exposed mice. As shown in **Fig. 3A**, the mean linear intercept in lungs was significantly increased by CS exposure, which was alleviated by digoxin administration (0.02 and 0.1 mg/kg). Goblet cell hypertrophy and hyperplasia can induce mucus hyper-secretion in the airway, which is one of the important factors of predicting the incidence and mortality of COPD (27). The PAS staining showed that compared with CTL group, the number of goblet cells in airway epithelium and the levels of mucin (Muc5ac and Muc5b) in BALF of CS group were significantly increased (**Fig. 3B**). Digoxin (0.02 and 0.1mg/kg) could effectively inhibit CS exposure-induced proliferation of airway goblet cells (**Fig. 3B**). But only high dose of digoxin (0.1mg/kg) can effectively inhibit the higher levels of Muc5ac and Muc5b in BALF induced by CS exposure (**Fig. 3C-D**). These results indicate that digoxin attenuates emphysema, goblet cell hypertrophy and hyperplasia, and mucus hyper-secretion in the airway of COPD mice.

### 3.4 Digoxin has no significant effect on liver, spleen and kidney indices and body weight of CS-exposed mice

CS exposure increased the kidney index (**Fig. 4C**) but decreased the body weight (**Fig. 4D**) of mice, and it has no effect on liver and spleen indices (**Fig. 4A-B**). Furthermore, digoxin (0.02 and 0.1 mg/kg) had no significant effect on the liver, spleen and kidney indices (**Fig. 4A-C**) and body weight of CS-exposed mice (**Fig. 4D)**. These results show that the doses of digoxin (0.02 and 0.1 mg/kg) used in this study have no apparent toxicity on important organs of mice.

**Figure 4.**
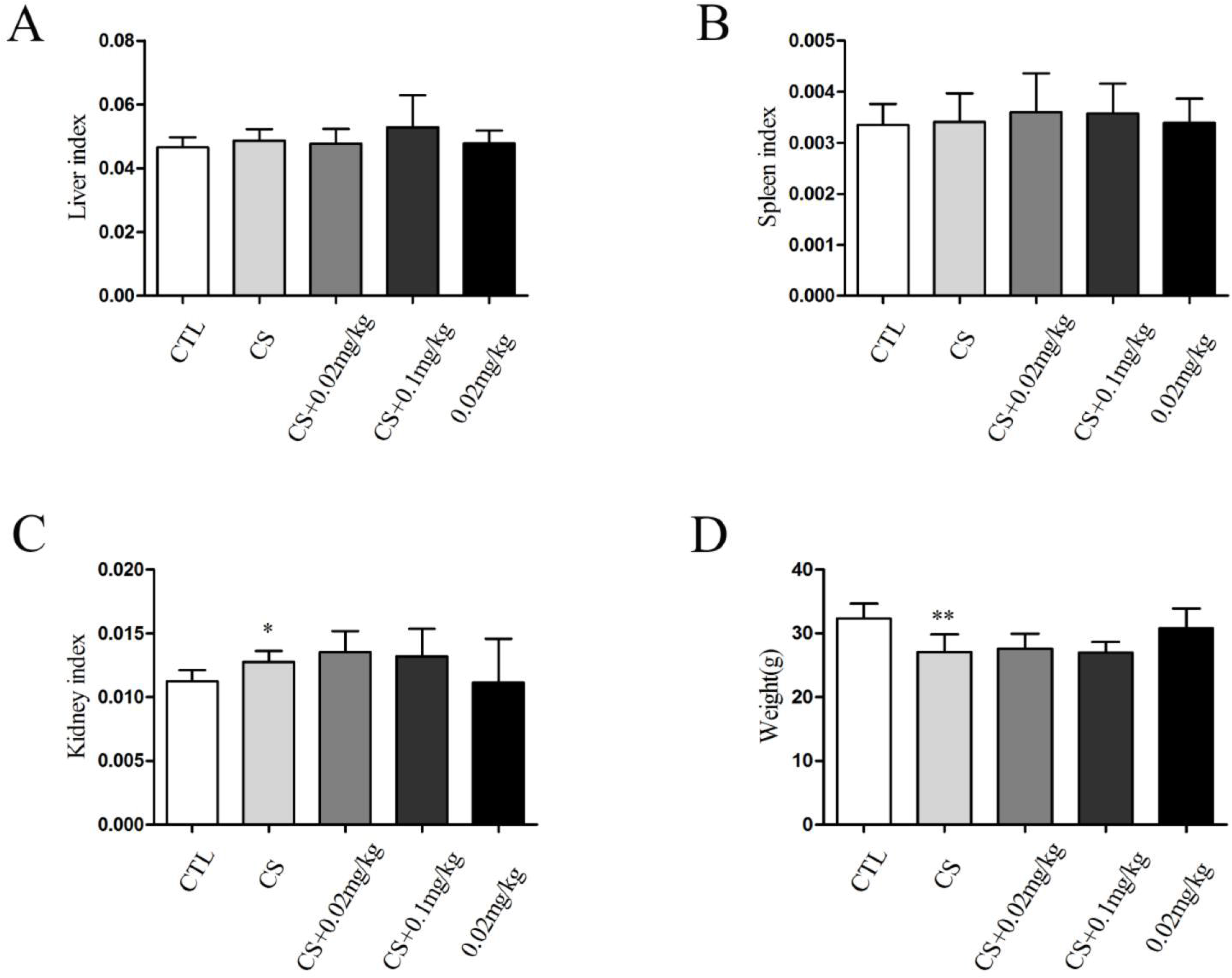
Digoxin has no significant effect on liver, spleen and kidney indices and body weight of CS-exposed mice. (A) liver index, (B) spleen index, (C) kidney index and (D) body weight of mice were measured. All data are shown as the mean ± SD; n= 8-10 per group in mice. Statistical significance was assessed by one-way ANOVA. ^**^*p< 0.01* versus control group.

### 3.5 Digoxin attenuates CS-induced or CSE-induced EMT in both mice and A549 cells

CS or CSE exposure can induce EMT *in vivo or vitro* (22, 28). Consistent with this, our results showed that CS exposure decreased the level of epithelial marker (Ecadherin) and increased the level of mesenchymal markers (Fibronectin and Vimentin) in mouse lungs (**Fig.5A**). Treatment with high concentration of digoxin (0.1mg/kg) significantly inhibited CS-induced EMT and HIF-1α.expression increase. Moreover, our *in vitro* data indicated that 2% CSE stimulation for 48 h significantly decreased the protein expression of E-cadherin and ZO-1, but increased the level of Vimentin and HIF-1α (**Fig.5B**). These changes were significantly inhibited by digoxin treatment in A549 cells (**Fig.5B**). These results show that HIF-1α contributes to CS-induced EMT both *in vivo* and *in vitro*.

**Figure 5.**
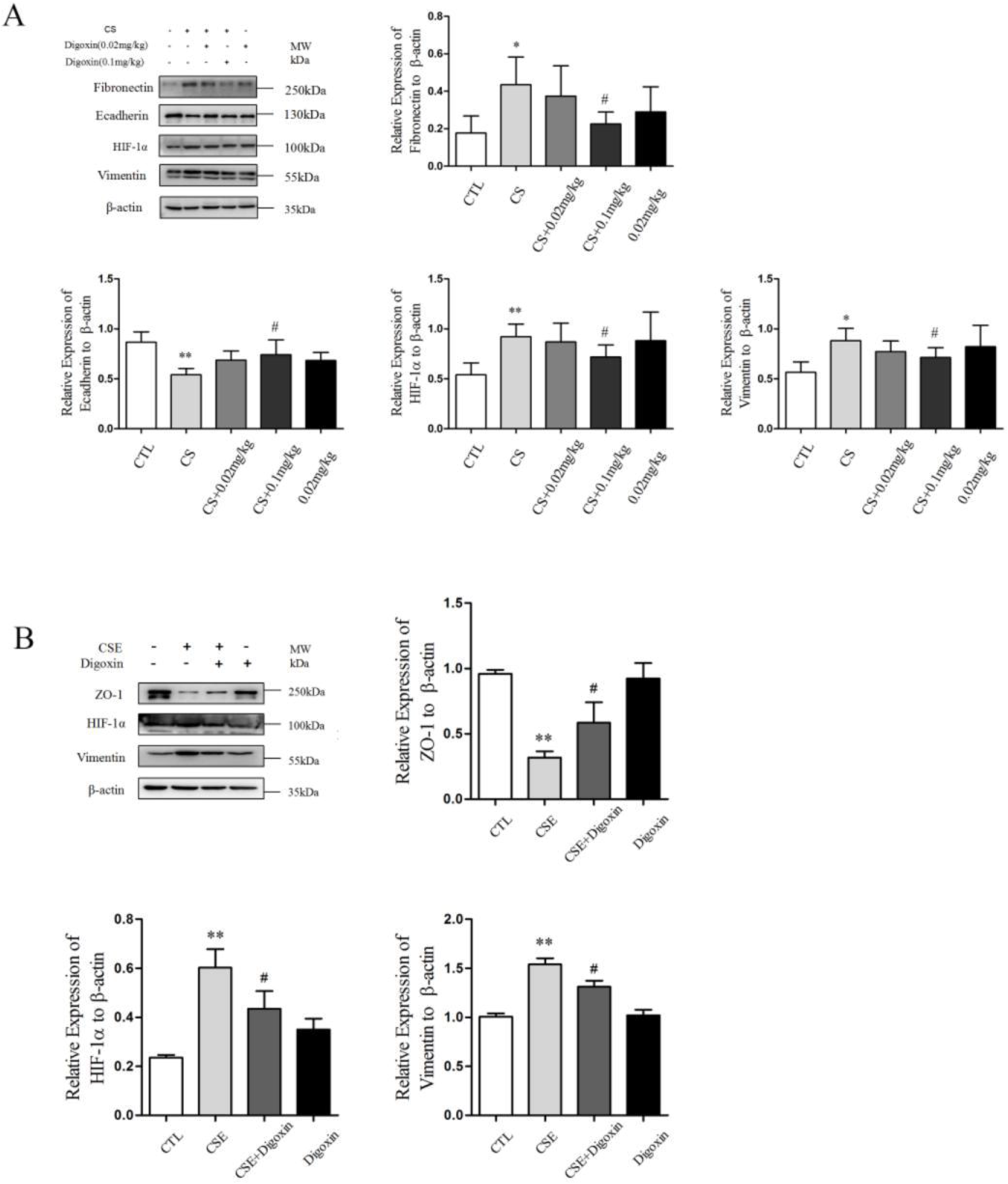
Digoxin attenuates CS-induced or CSE-induced EMT in both mice and A549 cells. (A) The expressions of Fibronectin, Ecadherin, HIF-1[, and vimentin in lung tissues were measured by Western blot. To further explore the effect of HIF-1α on CSE-induced EMT *in vitro*, A549 cells were treated with digoxin before 48-h CSE exposure. (B) The expressions of ZO-1, HIF-1[, and vimentin in A549 cells were measured by Western blot. All data are shown as the mean ± SD; n=8-10 per group in mice, n=5 per group in A549 cells. Statistical significance was assessed by one-way ANOVA. \**p< 0.05* and ^**^*p< 0.01* versus control group; ^#^*p< 0.05* and ^##^*p< 0.01* versus CS group.

### 3.6 Digoxin attenuates CS-induced or CSE-induced EMT possibly through HIF-1α /TGF-β1/Smad3 signaling pathway

TGF-β1/Smad signaling pathway plays a crucial role in triggering EMT (17), and HIF-1α protein synthesis is reported to regulate the TGF-β1/Smad signaling pathway (19). Therefore, we examined the effects of digoxin on HIF-1α, TGF-β1, and Smad mRNA levels both *in vivo* and *in vitro*. As **Figure. 6A-F** showed, CS or CSE exposure resulted in significant increase of HIF-1α, TGF-β1, and Smad mRNA levels both in mice and A549 cells, which was attenuated by digoxin treatment. To further explore the effect of TGF-β1/Smad pathway on CSE-induced EMT, we pretreated CSE-stimulated A549 cells with S7959 that can block the activation of Smad2/3. Pretreatment with S7959 inhibited the CSE-induced phosphorylation of Smad3 (**Fig. 6G**) and alleviated CSE-induced decrease of Ecadherin protein expression (**Fig. 6H**). These results indicate that digoxin suppresses CSE-induced EMT possibly through HIF1-α/TGF-β1/Smad pathway (**Fig. 6A-B**).

**Figure 6.**
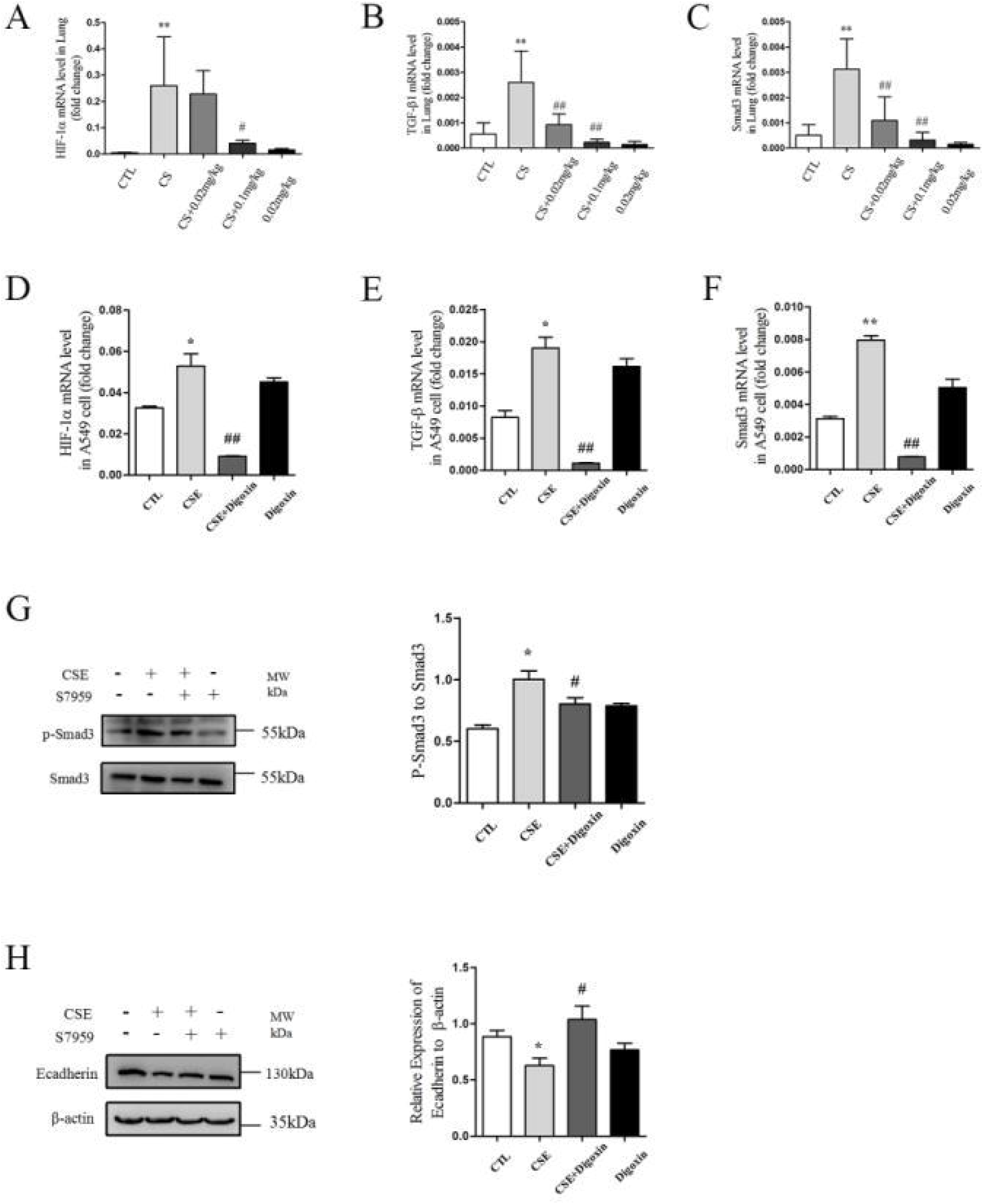
Digoxin attenuates CS-induced or CSE-induced EMT possibly through HIF-1α/TGF-β1/Smad3 signaling pathway. The mRNA levels of HIF-1α, TGF-β1 and Smad3 in lung tissues or A549 cells were measured by real-time PCR (A-F). To further determine the role of Smad3 in COPD-associated EMT, CSE-exposed A549 cells were pretreated with S7959 (a Smad3 inhibitor) for 1 h before 48-h CSE stimulation. The protein levels of p-Smad3 and E-cadherin in A549 cells were measured by Western Blot. All data are shown as the mean ± SD; n=8-10 per group in mice, n=5 per group in A549 cells. Statistical significance was assessed by one-way ANOVA. \**p< 0.05* and ^**^*p< 0.01* versus control group; ^#^*p< 0.05* and ^##^*p< 0.01* versus CS group.

## Discussion

HIF-1α is an important regulator of cellular responses to inflammation, oxidants, and hypoxia (5). Some studies have shown that HIF-1α may play a major role in COPD (6–9), however, its role in COPD has not fully investigated. In this study, we chose digoxin as the HIF-1α inhibitor to delineate the role of HIF-1α in the pathogenesis of COPD. Our study showed that digoxin, as a HIF-1α inhibitor, improved CS-induced inflammation and EMT both *in vivo* and *in vitro*. Moreover, digoxin alleviated CS-induced lung function decline, emphysema, and airway goblet cell hyperplasia in COPD mouse model. More importantly, our findings for the first time revealed that HIF-1α contributed to the development of EMT in CS-induced COPD possibly through TGF-β1/Smad3 signaling pathway.

CS, a major risk factor of COPD, can cause pulmonary inflammation, airway obstruction, and alveolar structure changes (29). In this study, COPD mouse model was induced by CS exposure for six months to simulate the development of COPD in human. At present, the clinical treatments of COPD are mainly bronchodilators and inhaled corticosteroids (ICS) (2). However, bronchodilators cannot slow down the decline of lung function and improve the prognosis (2), and ICS may increase the infection risk and acute exacerbation frequency of COPD patients (2). Moreover, EMT is implicated in respiratory structural remodeling and airway fibrosis in COPD and related to airflow obstruction (14), but no definitive treatment has been shown to effectively inhibit EMT in COPD (2). Therefore, it is urgent for us to find a new and effective method to prevent the development and improve the prognosis of COPD.

Our studied showed HIF-1α contributed to inflammatory responses, emphysema, goblet cell hypertrophy and proliferation in airway epithelium, airway mucus hypersecretion, and EMT in CS-induced COPD. It seems that HIF-1α is a promising intervention target of preventing the development and improving the prognosis of COPD.

The knockout of HIF-1α attenuates inflammatory responses of proximal colon cancer (30) and alleviates LPS-induced sepsis (31). Moreover, HIF-1α promotes hypoxia-induced inflammatory responses by activating the pathway of EGFR/PI3K/AKT (7). Consistent with these findings, our study showed that digoxin could alleviate CS-induced inflammatory cell infiltration and pro-inflammatory factor release in mouse lungs. At the same time, digoxin can significantly reduce CSE-induced secretion of IL-6 and IL-8 in A549 cells. However, the mechanism by which HIF-1α promotes CS-induced inflammation was not studied in depth. HIF-1α may inhibit the innate and adaptive immunity of airway epithelium (32), which will be focused in our future research.

Airway mucus hypersecretion, one of the important factors predicting the incidence and mortality of COPD (27), is closely related to airway hyper-reactivity (33) and airway obstruction (27). Studies have shown that controlling airway mucus hyper-secretion can effectively prevent acute exacerbation of COPD and improve quality of life of COPD patients (34). However, there is a lack of effective treatment for airway mucus hyper-secretion. Previous study has demonstrated that HIF-1α promotes hyperplasia of airway goblet cells via epidermal growth factor receptor-mediated signaling pathways (8). In addition, the binding of HIF-1α to conserved region of *MUC5AC* gene promoter has been confirmed, and it means that HIF-1α signaling pathway may be involved in up-regulation of airway mucus production (35). Our results also indicated that administration of digoxin inhibited airway goblet cell proliferation and mucus hyper-secretion, indicating that HIF-1α is a potential target for treating mucus hyper-secretion in COPD.

Pathological changes in COPD are characterized by emphysema, small airway remodeling and peribronchiolar fibrosis (2). EMT in airway epithelium is increased both in COPD patients and smokers with normal lung function (28). Our results also showed that CS exposure induced EMT both *in vivo* and *in vitro*. Moreover, EMT is involved in the respiratory structural remodeling and airway fibrosis in COPD (14), and contributes greatly to the occurrence and progression of COPD (10–13). Therefore, there is an enormous need to find new treatment targets for EMT in COPD. HIF-1α plays a key role in EMT of renal fibrosis and mammary cancer cells (15, 16). But so far, the role of HIF-1α on EMT in CS-induced COPD has not been reported. Our results suggested that administration of digoxin could significantly inhibit CS-induced increase of HIF-1α protein expression and alleviate CS-induced EMT in mouse lung tissues and A549 cells. CS exposure induces the formation of EMT mainly through Smad signaling pathways (22). Besides, HIF-1α can activate TGF-β1/Smad signaling pathway (19). This study also indicated that digoxin inhibited CS-induced increases of HIF-1α, TGFβ1, Smad3 mRNAs in mouse lung tissues and A549 cells. Moreover, the administration of Smad3 inhibitor was found to significantly inhibit the activation of Smad3 and CSE-induced EMT in A549 cells. Therefore, we believe that digoxin inhibits COPD-associated EMT through HIF-1α/TGF-β1/Smad3 signaling pathway. This demonstrates that HIF-1α may be a new therapeutic target for EMT in CS-induced COPD.

In summary, our study showed that digoxin as a HIF-1α inhibitor could significantly inhibit CS-induced emphysema, inflammatory responses, airway mucus hypersecretion, lung function decline and EMT. More importantly, we first found that digoxin could inhibit CS-induced EMT in COPD through HIF-1α/TGF-β1/Smad3 signaling pathway. This study provides a solid evidence for HIF-1α to contribute to the progression of COPD by promoting CS-induced EMT, inflammation, airway mucus hypersecretion and emphysema.

## Abbreviations

BALF: bronchial alveolar lavage fluid
COPD: chronic obstructive pulmonary disease
CS: cigarette smoke
CSE: cigarette smoke extract
EMT: epithelial-mesenchymal transition
MCP-1: monocyte chemoattractant protein-1
HIF-1α: hypoxia-inducible factor-1α

